# Age-related degradation of optic radiation white matter predicts visual, but not verbal executive functions

**DOI:** 10.1101/2020.11.04.368423

**Authors:** Christina E. Webb, Patricio M. Viera Perez, David A. Hoagey, Chen Gonen, Karen M. Rodrigue, Kristen M. Kennedy

## Abstract

Healthy aging is accompanied by degraded white matter connectivity, which has been suggested to contribute to cognitive dysfunction observed in aging, especially in relation to fluid measures of cognition. Prior research linking white matter microstructure and cognition, however, has largely been limited to major association and heteromodal white matter tracts. The optic radiations (OR), which transfer visual sensory-perceptual information from thalamic lateral geniculate nucleus to primary visual cortex, are generally considered lower-level input-relay white matter tracts. However, the role of this prominent white-matter visual relay system in supporting higher-order cognition is understudied, especially in regard to healthy aging. The present study used deterministic tractography to isolate OR fractional anisotropy (FA) in 130 participants aged 20-94 to assess age effects on OR tract white matter. We also examined associations between age-related differences in the OR and cognitive domains involving visual processing speed, and visual- and non-visual executive function (EF). OR microstructure, as indexed by FA, exhibited a significant linear decrease across age. A significant interaction between age, FA, and cognitive domain on cognitive task performance indicated that in older age, more degraded OR white matter was associated with poorer visual EF, but no age-related association between FA in the OR and visual processing speed or verbal EF was observed. These findings suggest the optic radiations are not merely sensory-perceptual relays, but also influence higher-order visual cognition differentially with age.

## Introduction

Typical aging is accompanied by degradation of structural white matter connections, which are essential for propagation of neuronal signals between gray matter networks. In particular, fractional anisotropy (FA), a measure of the degree of restricted diffusion, shows a negative linear association with age and is indicative of deterioration of microstructural organization. This age-related cortical disconnection is suggested to contribute to cognitive dysfunction observed in aging, especially in relation to measures of fluid cognition such as processing speed and executive function (1–5). In support of this theory, age-related performance variations in these cognitive domains are shown to be mediated by age differences in white matter properties (5–13). This evidence collectively demonstrates that age-related degradation of white matter connectivity contributes to poorer cognitive functioning in aging; yet, empirical evidence has largely been limited to the study of major association white matter tracts that link heteromodal gray matter regions supporting higher-order cognitive functioning. Because aging of white matter typically shows an anterior-to-posterior gradient (13–17), there has been a greater focus on linking age differences in structural properties of anterior white matter to age-related variation in cognition. There has been less research, however, specifically characterizing the extent of aging effects on posterior relay white matter pathways, such as the optic radiations (OR), that support vital sensory-perceptual processing. Moreover, it remains to be determined whether aging of this white matter influences not only performance on speeded tasks, but also more complex executive functions.

The optic radiations are large white matter bundles that originate from the lateral geniculate nucleus (LGN) of the thalamus and extend posteriorly to connect with the primary visual cortex (18), with a loop through the temporal cortex (i.e., Meyer’s loop). These radiations are responsible for the transfer of visual sensory-perceptual information and are generally thought to be lower-level input-relay tracts. White matter FA in the optic radiations has been linked to blood-oxygen-level-dependent (BOLD) activation in the visual cortex (19) and declines in OR FA can indicate visual impairment in older adults (20). While the OR facilitate transmission of visual information from the retina, this prominent white matter visual relay system also likely influences higher-order cognitive functions. However, the association between microstructural properties of the OR and their effect on processing speed or other complex cognitive functions remains largely understudied, especially in aging. In children, lower white matter FA in the OR tracts is associated with a significant reduction in processing speed (21), suggesting that reduced white matter quality of the OR results in decreased speed at which children process information and make decisions. Bells et al. (22) further showed that degraded OR white matter microstructure is related to neural synchronization in the visual cortex, which affects cognitive performance in children and adolescents. Additional evidence indicates that individual differences in younger adults’ performance on a choice reaction time task is associated with variability of FA in the OR, yet the sample size of this study was small (23). Only one study, to our knowledge, has specifically linked white matter properties of the OR with age-related differences in processing speed. Using principal component analysis (PCA), Johnson et al. (24) demonstrated that the OR were indeed sensitive to age-related differences in FA and that age mediated the association between FA and perceptual-motor speed. However, the specific relationship between OR white matter and its association with other higher-order visual and verbal cognitive functions has not been reported in healthy aging adults.

The present study sought to characterize the extent of age-related effects on FA in the OR in a lifespan sample of healthy adults. We also aimed to determine whether aging of OR white matter shows specificity in its association with visual versus non-visual, i.e., verbal, cognitive performance. We hypothesized that, consistent with other major white matter tracts, OR white matter FA would decrease linearly across the age span. Second, in line with the idea that the OR act as a visuospatial relay system supporting lower-level visual and perceptual functioning, we expected that aging of OR white matter would be related to basic visual processing speed. Age-related differences may also exist in the extent to which FA of the OR is associated with higher-order executive functions, and notably, may do so differentially across visual and verbal domains. Specifically, aging of the OR should influence performance on visual EF tasks to a greater degree than verbal EF tasks. To address these questions, the present study utilized diffusion tensor imaging (DTI) and deterministic tractography to isolate the OR and characterize both age effects and age-related differences in cognitive associations in a large lifespan sample of healthy adults.

## Methods

### Participants

Participants included 130 cognitively normal, healthy adults sampled from across the lifespan ranging in age from 20 – 94 years (mean age = 47.88, SD age = 17.36; 54 men, 76 women), who were recruited from the Dallas-Fort Worth Metroplex. Participants were screened against a history of cardiovascular, metabolic, or neurologic disease, head trauma with loss of consciousness, and any MRI contraindications, such as metallic implants and claustrophobia. Participants were screened before study entry for normal or corrected-to-normal vision (far and near vision ≤ 20/50). Including visual acuity (unit-weighted composite *z*-score of near and far acuity) as a covariate in final analyses did not change the pattern of results. Further inclusion parameters included a Mini-Mental State Exam (MMSE) (25) score of at least 26, and Center for Epidemiological Studies Depression(CES-D) (26) score of 16 or lower to exclude participants demonstrating indication of dementia or depression, respectively. **Table 1** reports participant demographics, as well as cognitive composite scores and mean OR FA, separated by arbitrary age group; all analyses treated age as a continuous variable. The study was approved by The University of Texas at Dallas and The University of Texas Southwestern Medical Center institutional review boards and all participants provided written informed consent.

**Table 1.**
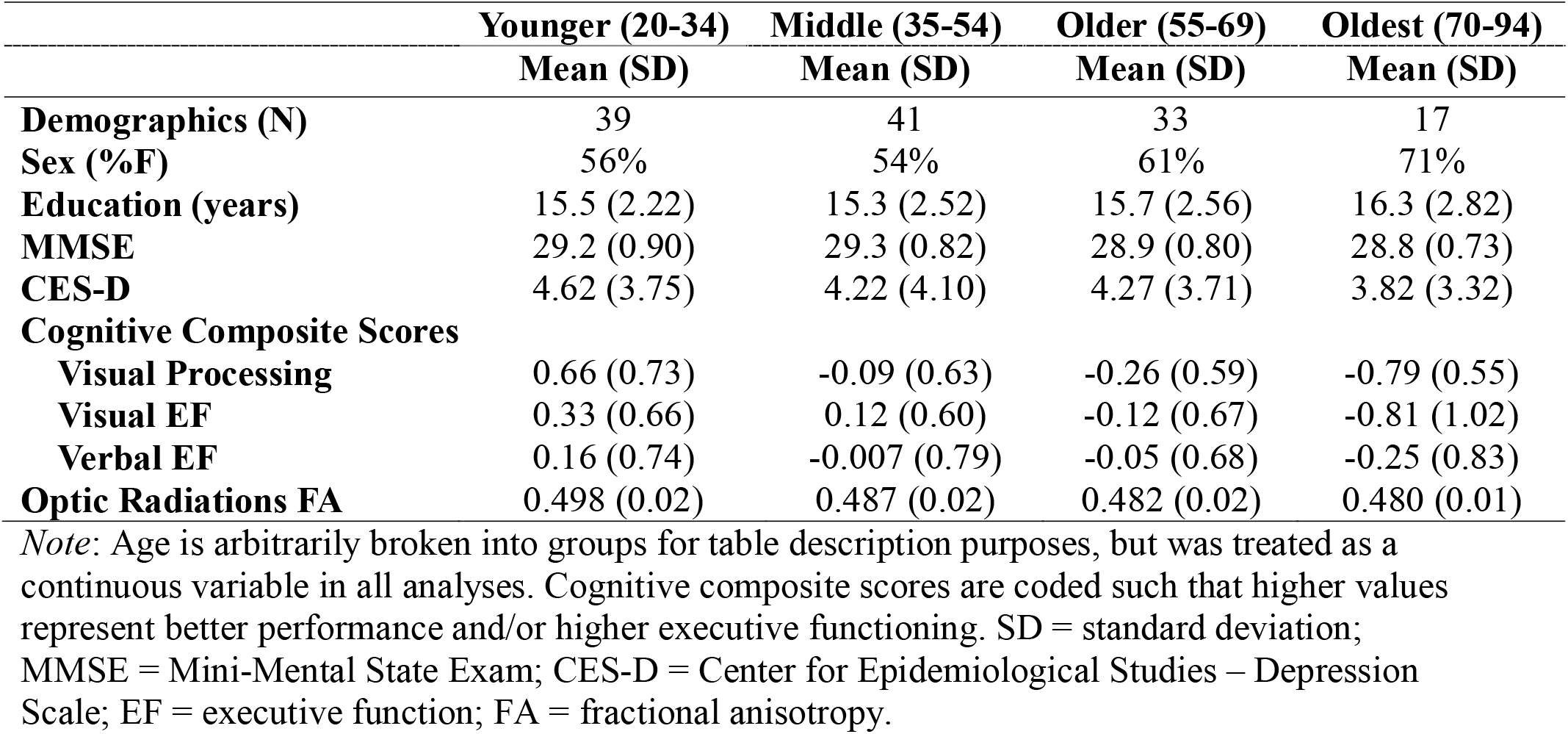
Participant demographics and cognitive performance

### MRI Acquisition Protocol

A 3T Philips Achieva MRI scanner equipped with a 32-channel head coil was used to acquire diffusion-weighted images with the following parameters: 65 whole brain axial slices in 30 diffusion-weighted directions (b-value = 1000s/mm^2^) with 1 non-diffusion weighted b_0_ (0s/mm^2^), voxel size 2 × 2 × 2.2 mm^3^ (reconstructed to 0.875 × 0.875 × 2 mm^3^), TR/TE = 5608 ms/51 ms, FOV = 224 × 224, matrix = 112 × 112. A high-resolution T1-weighted MPRAGE sequence was also acquired on the same scanner with the following parameters: 160 sagittal slices, voxel size 1 × 1 × 1 mm^3^, flip angle = 12°, TR/TE = 8.3 ms/3.8 ms, FOV = 256 × 204 × 160, matrix = 256 × 256.

### Diffusion Image Processing and Tractography Protocol

DTIPrep v1.2.4 was utilized for processing and quality control of diffusion images (27) and to identify potential susceptibility or eddy current artifacts. Gradients with intensity distortions, as well as those of insufficient quality caused by participant head motion, were detected and removed from subsequent analyses using the default thresholds in DTIPrep, and remaining gradients were co-registered to the non-diffusion-weighted b_0_ image. Diffusion directions were adjusted to account for independent rotations of any gradient relative to the original encoding direction (28). DSI studio (29) (software build from September 26th, 2014; http://dsi-studio.labsolver.org) was used to calculate the diffusion tensors and FA at each voxel and to conduct deterministic tractography of the OR. To ensure standardization across the sample, regions of interest and avoidance (ROIs; ROAs) were delineated on either the 1mm MNI template brain or by using the individual subject parcellations obtained through Freesurfer v5.3.0 (30) using the Desikan-Killany atlas (31). Regions were then warped to individual native subject diffusion space using a series of non-linear registrations that were calculated and applied using the Advanced Normalization Tools (ANTs) software package (32). ROIs included the left and right thalamus proper from Freesurfer segmentation, which was expanded to identify the white matter specific to the lateral geniculate nucleus (LGN), manually identified using the guidelines provided by Benjamin, et al. (33). Additionally, the calcarine sulcus was identified to extend tracking into the primary visual cortex (see **Figure 1**). The ROAs included a midsagittal plane so that tracking could be isolated to each hemisphere, and ROA planes located anterior (y = -2) and posterior/dorsal (y = -40) to the thalamus to exclude anterior/superior thalamic radiation projections or posterior thalamic radiation projections into parietal sensory association areas, respectively. Additional ROAs included the brainstem and cerebellum to prevent inferior projections. The following parameters were used in the deterministic tracking algorithm: maximum turning angle = 75 degrees, step size = 1 mm, minimum/maximum length = 20/500 mm, FA threshold = 0.20. Topology-informed pruning was used with 16 iterations to minimize wayward streamlines (34). Participants were required to have a minimum of 200 (1.25 SD from the mean) optic radiation streamlines across hemispheres. Lastly, mean FA (a scalar value ranging from 0 to 1) in the OR tract was calculated within hemisphere and then averaged across both hemispheres to obtain one mean OR FA value per participant. Mean global white matter FA (averaged from all white matter voxels in the brain) was also calculated to serve as a control variable in statistical models in order to account for any general influence of white matter FA on cognitive performance measures.

**Figure 1.**
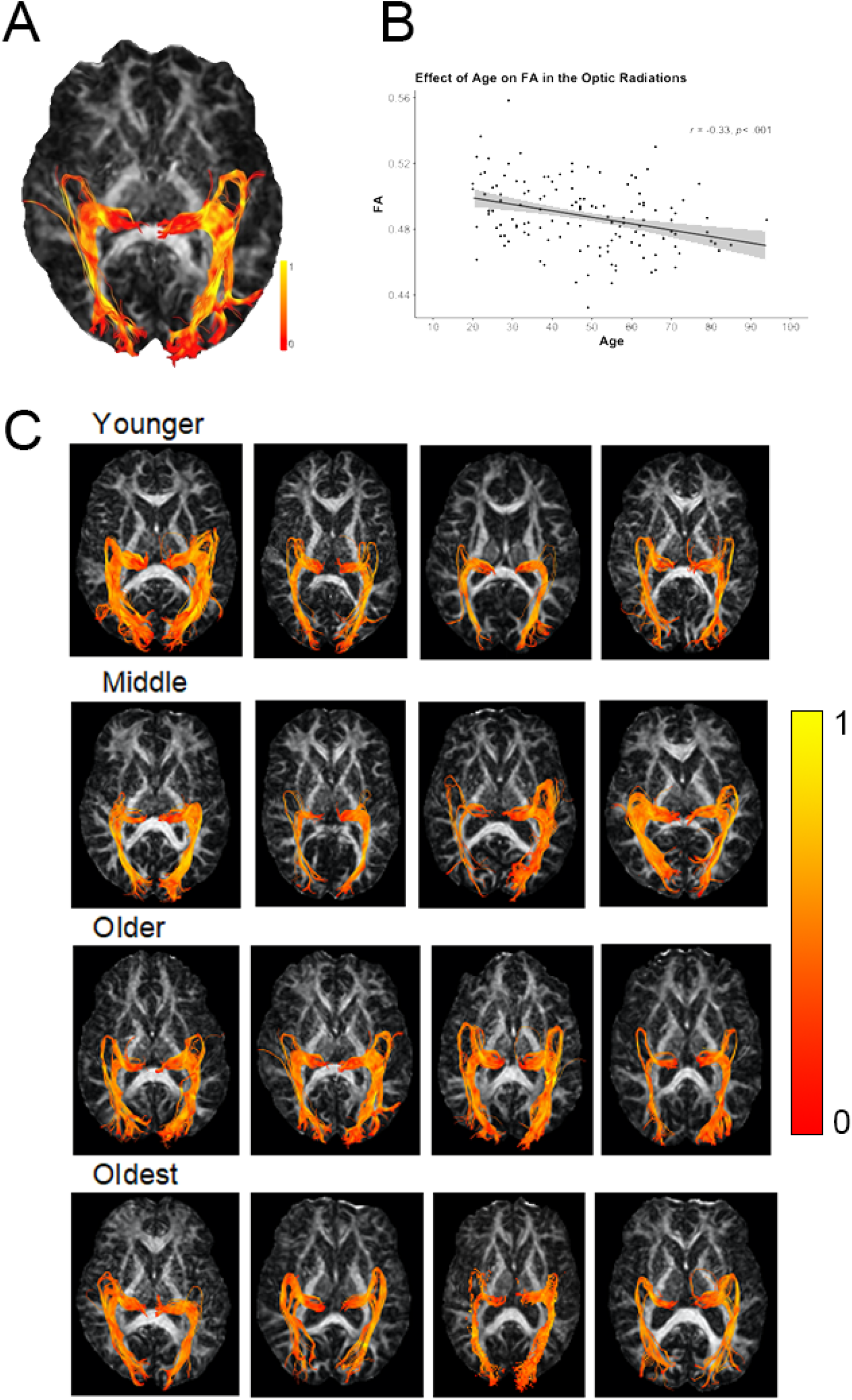
A) Axial view of the optic radiations (OR) tractography on a representative middle-aged participant’s fractional anisotropy (FA) map. Fibers extend from the lateral geniculate nucleus of the thalamus to the calcarine sulcus (V1 region). Meyer’s loop through the temporal cortex can also be seen. B) The scatterplot depicts the negative correlation between age and FA in the OR as well as illustrating the roughly equal distribution of participants across the adult agespan in this sample. Panel C) illustrates four random participants corresponding to each of the four lifespan “epochs” listed in Table 1. Heat gradient represents lower (red) to higher (yellow) FA values.

### Cognitive Measures

Three cognitive composites were created by averaging performance on cognitive tasks that reflect cognitive functions involving visual processing speed, visual EF, and verbal EF (detailed below). Performance on each cognitive task was coded so that higher scores reflected better performance (i.e., visual EF scores were multiplied by -1 to be reverse-coded), then unit-weighted composite *z*-scores were computed for each cognitive domain. All composites demonstrated a Cronbach’s α > 0.85.

### Visual Processing Speed

The visual processing speed composite was constructed from the Pattern Comparison Test and Letter Comparison Test, both of which require participants to make rapid judgements about whether or not two sets of stimuli are the same (35), as well as the Digit-Symbol Coding subtest of the Wechsler Adult Intelligence Scale (WAIS)(36), in which participants indicate associations between symbols and numbers as quickly as possible. All tasks capture aspects of visual processing speed, short-term visual memory, and visual-motor coordination. Performance on the Pattern and Letter Comparison tests (Parts 1 and 2) was calculated as the number of correct trials completed in 30 seconds for each part. Performance on the Digit-Symbol Coding test was calculated as the total number of correct trials completed in 120 seconds.

### Visual Executive Function

The visual executive function composite was created from the Color-Word Interference (Stroop) tests and the Trail Making Test as implemented in the Delis-Kaplan Executive Function System (D-KEFS) (37). The Color-Word Interference test involves color and word naming, inhibition, and cognitive switching conditions, and performance is measured as the total time taken (in seconds) to complete each condition. The Trails test involves visual scanning, number and letter sequencing, and cognitive switching conditions, and performance was measured as the total time taken (in seconds) to complete each condition. Scores on the visual EF tasks were reverse-coded, so that higher scores reflect better EF.

### Verbal Executive Function

The verbal executive function composite was formed from the D-KEFS Verbal Fluency test, which assesses an individual’s ability to generate words fluently in effortful, phonemic format, from general categories, as well as a condition involving shifting between categories. Performance was measured as the total number of words correctly produced in each condition (including F, A, S, Animals, Boys, and switching conditions) corrected for any set loss or repetition errors. Thus, the verbal EF composite was constructed of six verbal fluency subtest scores.

### Statistical Analysis Approach

A linear regression was used to evaluate the effect of age on OR FA. Linear mixed-effects models were conducted to estimate whether age and OR FA are differentially related to performance on the visual and verbal cognitive measures. Specifically, a model including between-subjects fixed effects of continuous FA and continuous age, a within-subject fixed effect of cognitive domain (visual processing speed, visual EF, verbal EF), and a random intercept on participant, as well as interactions between FA, age, and cognitive domain, was estimated predicting cognitive performance. Modeling both within- and between-subject effects in this way allows for proper estimation of standard errors when observations are not independent (e.g., cognitive performance across domains within an individual). To account for potential sex differences and effects of education on the measures of cognition, the model also included covariates of sex and education. To mitigate the likelihood that effects were representative of general white matter effects, the model additionally included a covariate representing global, or whole brain FA. Interactions between all covariates and the within-subject fixed effect of cognitive domain were also included. All analyses were conducted using the “*lm*” and “*lmer*” functions implemented in R statistical software program (38), and in all models continuous variables were mean-centered to minimize multicollinearity. Significant interaction terms between continuous variables were followed-up with simple slopes analyses using the Johnson-Neyman procedure (39) on estimates from the mixed-effects models using the “*interactions*” package in R. This procedure allows for estimation of regions of significance of the effect of the independent variable on the dependent variable across values of the continuous moderator (40,41). In the current context, it allows for the pinpointing of when in the age span FA in the OR is significantly related to cognitive performance within each domain (i.e., at what value of age in years is the slope of the FA-cognition association significant and/or insignificant).

## Results

### Effect of Age on Optic Radiations

There was a significant negative effect of age on white matter FA in the optic radiations, where, as age increased FA decreased linearly [*F*(1,128) = 15.54, *p* < .001; see **Figure 1**].

### Age-related Effect of Optic Radiations on Cognitive Measures

Linear mixed-effects models were conducted to determine differential age effects of OR FA on performance across the cognitive domains. A conditional effect of age on cognitive performance was observed [*F*(1,123) = 36.48, *p* < .001] with performance decreasing with increasing age. There was also a significant age by cognitive domain interaction on performance [*F*(2,246) = 16.98, *p* < .001]; however, this was qualified by the presence of a three-way interaction among age, FA, and cognitive domain [*F*(2,246) = 3.30, *p* < .05]. No other model effects were significant (*p*s > .09). Decomposition of the three-way interaction indicated that there was a significant difference in how aging influences the FA-visual EF association versus how it influences the FA-visual processing association [*Est(SE)* = -0.45(0.19), *t*(246) = -2.36, *p* < .05; b = -.45 (95% CI: [-0.764, -0.136], SE = 0.19, t(246) = -2.36] and FA-verbal EF association [*Est(SE)* = -0.39(0.19), *t*(246) = -2.05, *p* < .05; b = -.39 (95% CI: [-0.704, -0.076], SE = 0.19, t(246) = -2.05]. No significant difference in age moderation of the FA-visual processing versus FA-verbal EF associations were detected [*Est(SE)* = 0.06(0.19), *t*(246) = 0.32, *p =* .75]. Simple slopes analyses, which allow the assessment of the nature of the association between FA and cognition at varying points in the age span, indicated that higher FA in the optic radiations was significantly associated with better performance on visual EF tasks in older age (beginning at about age 58; **Figure 2**). While the simple slopes analysis suggests that this age association flips in direction prior to about age 21, this approaches the boundary of the lower range of age in our data, and thus is less reliable and will not be further discussed. Further research including individuals earlier in the age span, such as adolescents, is needed to investigate whether there are different developmental trends in the association between visual white matter microstructure and higher-order cognition. Importantly, simple slopes analyses indicated no significant association between OR FA and performance on the visual processing or verbal EF tasks at any point in the age span, demonstrating cognitive domain specificity^1^.

**Figure 2.**
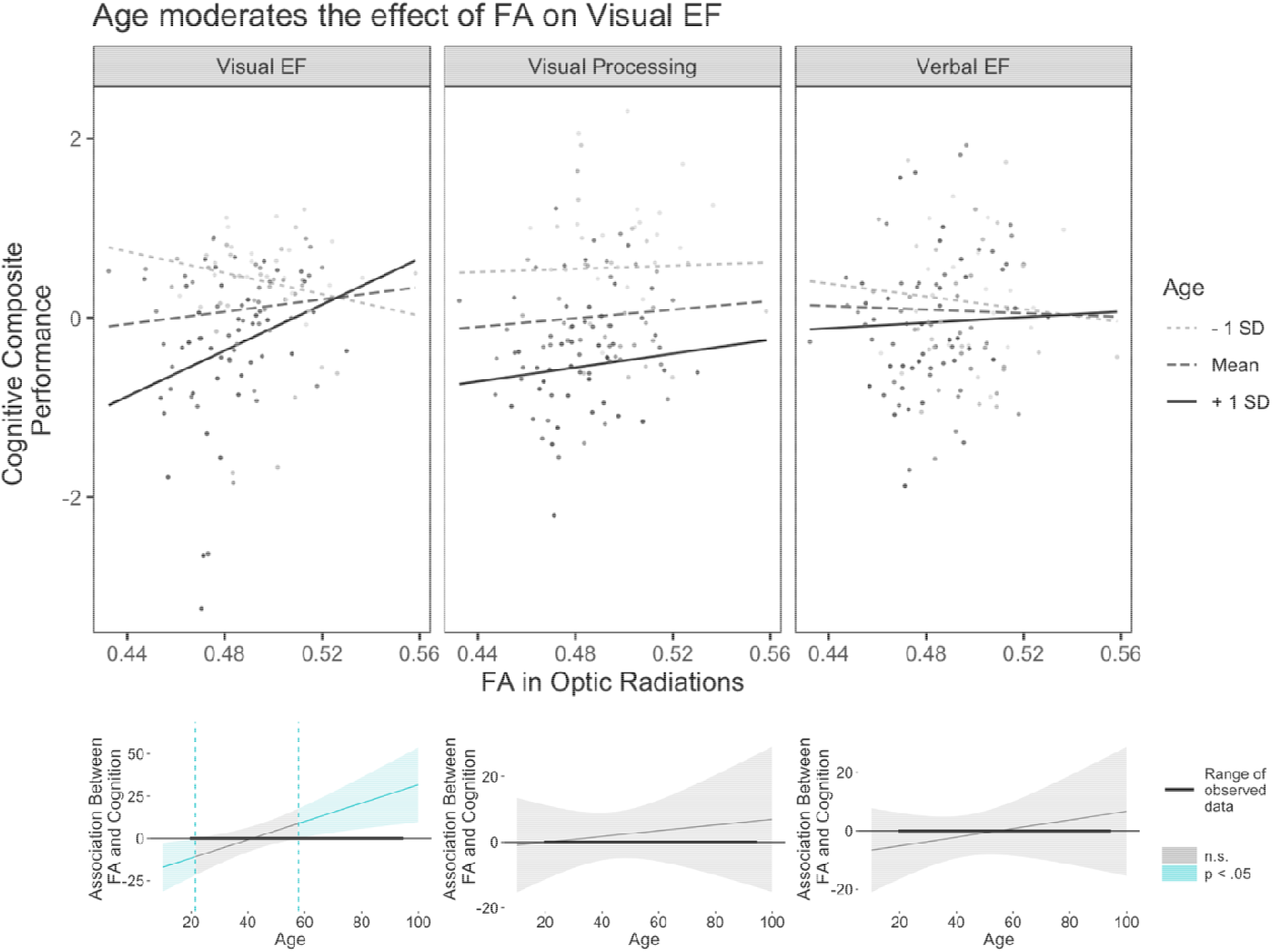
Age moderates the association between fractional anisotropy (FA) in the optic radiations (OR) and cognitive performance differentially across domain. For visual executive function (EF), higher FA in the OR is associated with better EF only in older age (from about age 58 onward), while there was no significant differential association between FA and visual processing or verbal EF across age. In the top set of graphs, slopes are depicted at -1 standard deviation (SD), mean, and +1 SD of age (age 30, 48, 65, respectively). The FA-visual EF performance association is significant for +1 SD of age (*t* = 2.36, *p* < .05), but non-significant (n.s.) at -1 SD (*t* = -1.36, *p =* .17) and at the mean of age (*t* = 0.93, *p =* .35). Individual data points depict observed data. The bottom set of graphs depict regions of significance plots for each cognitive domain indicating at what values of age the association between FA in the OR and cognition is significant (blue shaded regions) and non-significant (gray shaded regions); left: visual EF, middle: visual processing speed, right: verbal EF.

## Discussion

The present study utilized deterministic tractography to isolate the optic radiations, posterior white matter tracts essential for transmission of visual-sensory information, in a large lifespan sample of healthy adults. The visual cortex, along with other sensory gray matter regions, appear to be relatively resistant to the effects of aging compared to other brain regions (e.g., 40,42), yet to date, sparce research on aging of white matter projection tracts innervating the visual cortex is available. Here, we demonstrate that FA in the OR decreases linearly across the lifespan, contributing novel evidence to current literature showing age-related degradation of major association and commissural white matter tracts in normative aging. These age differences in FA were related to individual variation in cognition differentially across visual and verbal cognitive domains. While FA was unrelated to performance on a visual processing speed composite or to verbal fluency, age-related degradation of white matter in the OR was associated with poorer performance on tasks measuring visual executive functioning. Results contribute to the extant literature by characterizing age-related differences in microstructural properties of the OR and demonstrating effects of diminished visual white matter on executive function in healthy cognitive aging.

### Age Effects on Optic Radiation White Matter

A wealth of previous research indicates that white matter connectivity throughout the brain decreases with aging, with regional variation in the strength of the age association (13–16). Posterior white matter, especially in the occipital lobe, displays earlier maturation and declines at a later age compared to frontal areas, exhibiting a reversal of developmental myelin maturation (2,17,43). However, there has been a lack of studies specifically illustrating the extent of aging of OR white matter. Those that have included visual white matter have utilized ROI-based or voxelwise approaches, making it difficult to delineate age effects on specific white matter bundles of the visual system. Here we utilized a deterministic tractography approach to isolate the OR projection tracts and examine age differences across the adult lifespan. Results indicated that OR white matter FA decreased linearly across the age span, consistent with previous research detailing age trends of white matter tract FA throughout the brain (11,44). While FA is an indirect measure of white matter health and the exact property of white matter it captures is debated (45), age-related degradation of FA is generally thought to reflect degeneration of myelinated fibers in white matter (46). Our results suggest that white matter FA in the OR is susceptible to age-related alteration in microstructural organization, exhibiting a linear decrease across the lifespan. Future studies including longitudinal measures of white matter microstructure of the OR are required to determine the extent to which OR white matter properties change over time, and to identify to what degree the magnitude and trajectory of age-related decline in the OR compares to that of other association and projection fibers.

### Age-related Effect of OR White Matter on Cognition

The optic radiations act as a relay system connecting the lateral geniculate nuclei of the thalamus to primary visual cortices and facilitate processing and integration of sensory-perceptual information. Age-related degeneration of white matter in this region should consequently influence functional brain processes involving low-level information processing and/or higher-order cognitive domains. Our results indicate that aging of OR FA is associated with cognition differentially across visual and verbal aspects of cognition. Across the lifespan, OR FA showed little association with performance on tasks measuring speed of visual information processing or with those assessing verbal EF. Instead, age moderation of the association between FA in the OR and visual EF was observed, with negative effects of decreased white matter on visual EF prevalent beginning in late-middle age and continuing into older age (at about age 60 and above). Thus, possessing more degraded OR white matter in older age is associated with exhibiting poorer performance on visual EF tasks, supporting existing theories of cortical disconnection and white matter degeneration in aging (1,2,3,44).

White matter FA has been previously linked with individual variation in performance on tasks assessing processing speed in aging populations (1,3,4,47). Several studies also report that white matter FA in the frontal and parietal cortices (e.g., genu and splenium), as well as whole-brain FA, mediates age-related reductions in speed of processing (9,48,49). In regard to the OR specifically, previous studies demonstrate that reductions in the structural properties of OR tracts are associated with decreased processing speed in children and younger adults (21–23); yet, another study found no association between any OR diffusion metric measured and simple decision/reaction time in younger adults (50). The association of the OR tract with processing speed or other higher-order cognitive functions, however, has not been thoroughly investigated in a healthy aging adult population. In the present study, FA in the OR was not associated with simple visual processing speed at any point in the agespan, contrary to our initial predictions. While one other study including both younger and older adult groups found that age-related decreases in OR FA were associated with slower processing speed (24), mediation analyses indicated that age accounted for the majority of the variance between FA and the speed measure, suggesting that any association between FA and processing speed may be a mere reflection of the aging process. In the present study, when processing speed is considered independently in a separate linear model predicting OR FA, there is a significant positive association between speed and white matter FA; however, this association does not remain after accounting for effects of age and/or global FA on processing speed. Thus, together with our results, this evidence suggests that although the OR are thought to play an important role in transfer of information from the visual cortices, OR white matter structure is not distinctly related to processing speed beyond the effect of age.

Instead, our results indicated that FA in the OR was associated with significantly poorer performance on EF tasks. This association was modality specific, with an age-dependent FA effect exclusively observed for the visual, but not verbal EF composite. Notably, the positive association between OR FA and visual EF was evident only in late middle to older age (> about age 60), supporting a white matter connection degeneration hypothesis of aging. Numerous studies have demonstrated that age-related decline in white matter microstructural organization is associated with EF across various association and projection white matter tracts (3). Despite this evidence, a dearth of research exists investigating how aging of posterior white matter that supports information relay in the visual system contributes to EF. In support of the present results, Kennedy & Raz (5) found that reduced FA in posterior white matter (specifically ROIs in parietal, splenium, and occipital white matter) was associated with poorer performance on visual EF tasks measuring inhibition (i.e., Stroop) and task-switching. The present study contributes novel evidence specifically linking white matter microstructural properties of the OR to higher-order cognition. Critically, this OR structure-cognition association was observed after accounting for the general influence of white matter (i.e., global FA) on visual EF, providing evidence that individual white matter tracts contribute to complex visual cognition beyond a global effect of white matter health (24). Our results suggest that diminished white matter connectivity of the OR appears even during the course of healthy aging and is, in part, responsible for poorer visual executive performance that is characteristic of aging.

The fact that white matter in the OR had a specific age-dependent effect on visual, but not verbal EF, is in line with its established role in supporting the transfer of visual information from the lateral geniculate nucleus (originating from the retina) to the visual cortex. This dissociation extends previous structure-function studies which often combine visual and verbal EF measures, making it difficult to distinguish the relative influence of white matter degradation on age-related variation in executive performance between domains. Future research should evaluate how aging of different white matter tracts differentially relates to unique components of EF and other cognitive processes, such as working memory. While it was expected that FA in the OR would also be related to speed of visual information processing, it may be the case that the synchronization required for higher-order visual functioning is impacted first by OR degradation, and performance on speeded tasks is not linked to FA in the OR until later in the aging or disease process. Testing this hypothesis, however, requires longitudinal measurement of the rate of change in structure-function associations, currently underway in our lab. Alternatively, age-related degradation of white matter may influence speed of information processing in a more global manner and may not be isolated to particular association or projection tracts. Regardless, the present results suggest that OR white matter microstructure is involved in facilitating aspects of higher-order functions that rely on visual information (e.g., cognitive switching, visual-motor sequencing), rather than simply being a visual sensation and perception relay tract, and moreover, that age-related degradation of this white matter system is accompanied by poorer visual cognition.

Results from this study should be considered in light of various strengths and limitations. Inclusion of a sample spanning the adult lifespan, as opposed to younger and older age groups, allows for investigation of continuous age-related trends in how white matter microstructure is associated with individual differences in cognition across domains. Utilization of deterministic tractography to isolate individual OR white matter tracts as opposed to sampling from regions of interest placed within a portion of the OR, or measuring only a subsection of the OR through voxel-wise approaches, allows for a better representation of OR microstructural properties (51,52). Through a mixed-model approach we were also able to simultaneously compare the relative association between age-related alterations in OR white matter and performance across several cognitive domains versus examining these associations independently. Relatedly, while a strength of the present study was the inclusion of multiple measure of cognition in each domain, additional research using alternative speed measures could potentially show an association between FA in the OR and processing speed akin to that seen in other association white matter tracts. As with any study quantifying properties of white matter, FA represents an indirect property of white matter, and the true biological processes of white matter that it characterizes is a topic of continued debate. In addition, the health of white matter is only one biological property that differs across age, and it is likely that a multitude of factors (e.g., cortical thickness, gray matter volume, brain iron or beta-amyloid levels) contribute to age-related decline in cognition. Longitudinal data using multimodal neuroimaging methods are necessary to quantify individual age-related change in OR white matter degradation, as well as to identify associations between micro and macrostructural properties and cognitive performance across time.

Overall, the present study demonstrates that degraded white matter connections between thalamic regions and visual cortices occurs in a linear fashion across the lifespan. This weakened microstructure of the optic radiations is also linked to poorer cognitive performance, demonstrating the greatest effect on visual components of executive functioning in advanced age. Results extend the limited previous work characterizing structure-function associations to posterior white matter of the visual relay system and lend further support to theories suggesting that cortical disconnection contributes to declines in cognition observed in normal aging.

## Data availability statement

The data and relevant analysis code will be made publicly available via the Open Science Framework upon publication.

## Footnotes

1 Our *a priori* metric of interest was FA; however we observed a similar interaction and interaction breakdown effect for Visual EF when other diffusion metrics were considered (i.e., for MD and RD but not AD).

